# Cross-modal orienting of exogenous attention results in visual-cortical facilitation, not suppression

**DOI:** 10.1101/2020.10.13.338210

**Authors:** Jonathan M. Keefe, Emilia Pokta, Viola S. Störmer

**Affiliations:** Department of Psychology, University of California, San Diego, 92092, USA; Department of Brain and Psychological Sciences Dartmouth College

## Abstract

Attention may be oriented exogenously (i.e., involuntarily) to the location of salient stimuli, resulting in improved perception. However, it is unknown whether exogenous attention improves perception by facilitating processing of attended information, suppressing processing of unattended information, or both. To test this question, we measured behavioral performance and cue-elicited neural changes in the electroencephalogram as participants (N = 19) performed a task in which a spatially non-predictive auditory cue preceded a visual target. Critically, this cue was either presented at a peripheral target location or from the center of the screen, allowing us to isolate spatially specific attentional activity. We find that both behavior and attention-mediated changes in visual-cortical activity are enhanced at the location of a cue prior to the onset of a target, but that behavior and neural activity at an unattended target location are equivalent to that following a central cue that does not direct attention (i.e., baseline). These results suggest that exogenous attention operates solely via facilitation of information at an attended location.

## Introduction

In order to deal with the surfeit of information entering our senses, we can engage spatial selective attention voluntarily based on current goals or orient attention reflexively to a salient event in the environment, which both result in improved perception at the attended location (Wright & Ward, 2008; Carrasco, 2011). Foundational theories of attention propose that two mechanisms are involved during spatial selection: enhancement of sensory processing at the attended location and suppression of sensory processing at the unattended location (Desimone & Duncan, 1995; Pinsk, Doniger, & Kastner, 2004). This is based on the idea that visual processing is a strictly limited resource, and that if visual processing is increased at one location then it should be paralleled by a decrease in visual processing elsewhere. Studies of endogenous (i.e., voluntary) spatial attention have used electroencephalography (EEG) to examine neural activity following spatial attention cues, and found that when a location is attended, neural activity in parts of the visual cortex that represent that location is increased relative to other unattended locations even before a target stimulus is presented (e.g., Nobre et al., 2000; Grent-‘t-Jong & Woldorff, 2007). Several studies have argued that these preparatory biasing signals not only reflect neural enhancement of attended regions, but also anticipatory suppression of unattended areas (Serences et al., 2004; Kelly et al., 2006; McDonald & Green, 2010; Couperus & Mangun, 2010; Snyder & Foxe, 2010). More recently, a similar preparatory effect on visual-cortical processing was observed during the exogenous orienting of attention. Following peripheral salient sounds that oriented attention to the left or right side of space, neural activity was increased over the occipital cortex contralateral to the sound’s location relative to ipsilateral, even prior to or in absence of visual targets (McDonald et al., 2013). However, based on this relative difference in neural activity between the hemispheres, it cannot be distinguished whether this modulation reflects signal enhancement at the attended location, suppression of the unattended location, or a combination of the two. Thus, while many studies have attempted to tease apart mechanisms of enhancement and suppression during endogenous attentional orienting, research on how exogenous attention modulates neural activity in anticipation of a target is lacking.

We addressed this issue by adapting a cross-modal cueing paradigm to include a baseline ‘no-shift’ cue that allowed us to isolate neural activity related to the attended and unattended locations. Prior behavioral studies have used such baseline cues to separate behavioral benefits and costs of target processing, finding evidence for both attentional benefits (i.e., increases in performance for the cued location relative to baseline; e.g., Yeshurun & Carrasco, 1998; Carrasco, Penpeci-Talgar, & Eckstein, 1999) as well as attentional costs (i.e., decreases in performance at the uncued location relative to baseline; Quinlan & Bailey, 1995; Montagna et al., 2009; Pestilli & Carrasco, 2005). However, based on behavioral responses alone it is difficult to infer how orienting spatial attention is implemented in sensory cortex, given that responses are the result of multiple processing stages including preparatory sensory activity, perceptual processing of the target, and subsequent cognitive stages that involve decision-making, response preparation and response execution. Thus, in addition to behavior, we here directly assessed visual-cortical processing during exogenous attentional orienting using EEG.

Participants performed a visual discrimination task, and shortly before a visual target appeared, an auditory cue was presented either at the left or right target location to orient spatial attention, or at a central location – in this case acting as an alerting signal without eliciting lateralized shifts of attention (no-shift cue). Note that we used sounds as attention cues to eliminate any early visual responses elicited by the cues themselves. Our EEG analysis focused on two event-related potentials (ERPs). First, we examined the Shift-Related Positivity (SRP) over frontal electrode sites to confirm that our no-shift cue differed from the shift-cues in terms of engaging attentional control areas (Störmer, Green, & McDonald, 2009); second, we examined the Auditory-Evoked Contralateral Occipital Positivity (ACOP), a positive deflection over contralateral relative to ipsilateral occipital cortex with respect to the cue location (McDonald et al., 2013; Keefe & Störmer, 2020). The main question was how these lateralized changes over occipital cortex would compare to activity elicited by the no-shift baseline cues. Specifically, if exogenous attention is primarily supported by neural enhancement, we would expect increased activity over contralateral cortex relative to the no-shift cue; alternatively, if exogenous attention is primarily supported by neural suppression, we would expect decreased activity over ipsilateral cortex relative to the central no-shift cue; finally, an intermediate level of activity elicited by the no-shift cue would suggest the involvement of both enhancement and suppression.

## Method

### Participants

Nineteen participants were included in the final sample of the experiment (14 female; mean age of 19.9 years). Data from five participants were excluded due to excessive artifacts in the EEG (affecting > 30% of trials). Data from two additional participants were excluded due to issues with the EEG system that resulted in significant lost data: an HEOG electrode came loose for one subject and a battery died for the other. Two subjects lost a small number of trials (11 and 14 trials) due to sampling errors of the EEG system but are included here, as each subject had greater than 70% of the full sample trial number remaining following artifact rejection.

All participants gave informed written consent as approved by the Human Research Protections Program of the University of California, San Diego and were paid for their time ($10/hour) or received course credit. All participants reported having normal or corrected-to-normal vision and normal hearing. The sample size was chosen a priori based upon a number of other studies utilizing similar cross-modal attentional cueing paradigms that effectively measured the ACOP and/or related behavioral effects (McDonald, Teder-Salejarvi, & Hillyard, 2000; Störmer, McDonald, & Hillyard, 2009; McDonald et al., 2013; Feng et al., 2014; Keefe & Störmer, 2020). We preregistered our predictions and analysis on AsPredicted and planned to collect data from 20 participants after exclusion. Our sample has one fewer subject than planned due to data collection being disrupted by COVID-19.

### Stimuli and Apparatus

Participants were seated approximately 45 cm in front of a 27” monitor in a sound-attenuated, electrically shielded booth. Stimuli were presented on the screen via the Psychophysics Toolbox in MATLAB (Brainard, 1997; Pelli, 1997). A small black fixation dot (0.2° × 0.2° of visual angle) was always present in the center of the screen, which was otherwise uniformly gray (RGB: 127, 127, 127). A black circle (0.4° × 0.4°) appeared around the fixation dot at the start of each trial to indicate to the participant that the trial had begun. Peripheral auditory cues were ~83 ms pink noise bursts (0.5–15 kHz, 78 dB SPL) played from external speakers mounted on either side of the computer monitor. These cues were played in stereo and their amplitude was adjusted to give the impression that the sounds were emanating from the possible target locations on the screen (i.e., for a left cue, the amplitude of the sound coming from the left speaker was adjusted to be louder than the amplitude of the sound coming from right speaker, so that the sound appeared to come from the exact location of the visual target location). The central auditory cue was the same pink noise burst played from a speaker mounted on the top of the monitor (hereafter referred to as *central cue*) and adjusted to be equal in intensity to the peripheral stimuli. The central cue appeared to emanate from the center of the screen, and thus would not elicit any lateralized shifts of spatial attention. The target was a Gabor patch with a spatial frequency of 1.3 cycles/degree, turned either −45° or 45° from vertical. The contrast of the Gabor patch was determined for each participant in a calibration task prior to the main experiment (see below). The target was presented in one of two peripheral locations indicated by a black circle with a diameter of ~9° visual angle, centered ~28° of visual angle to the left and right of fixation. Each target was immediately followed by a visual noise mask of the same size.

### Procedures

Participants were asked to keep their eyes on the central fixation dot throughout each experimental block. Each trial began with the presentation of a black circle that appeared around the central fixation dot after 500 ms, indicating to the participants that the trial had started.

Following the onset of this circle at a variable inter-stimulus interval (ISI) of 500 – 800 ms, an 83-ms auditory attention cue was presented randomly at either the left, right, or center and was not predictive of the spatial location of the visual target. Consequently, participants were instructed to ignore the sounds because they would not be informative to the task. On 1/3 of trials, a target was presented following the cue at a stimulus onset asynchrony (SOA) of 130 ms, and on another 1/3 of trials the target was presented following the cue after 630 ms. The target Gabor patch was presented at one of the two peripheral locations for ~50 ms, and was followed immediately by a visual noise mask for 100 ms. The noise mask always appeared at the location of the target to eliminate uncertainty about the location at which the target appeared. Following the noise mask at an ISI of 300 ms, the black circle surrounding the central fixation dot turned white, prompting a response from the participant as to which direction the target was oriented. Participants made this report using the “m” (clockwise) and “n” (counterclockwise) keys. On the remaining 1/3^rd^ of trials, no target appeared and no visual information was presented following the cue. These trials ended 900 ms after cue onset, and participants pressed the spacebar to continue to the next trial. The target display was omitted or presented after a longer cue-target SOA (630ms) to allow recording of the event-related potential (ERP) to the cue separately from the otherwise overlapping ERP to the visual target.

All trial types were randomly intermixed within a block. Subjects performed 12 blocks of 72 trials each. Prior to the experimental tasks, task difficulty was adjusted for each participant using a thresholding procedure that varied the contrast of the Gabor patch target to achieve about 75% accuracy (i.e., QUEST; Watson & Pelli, 1983). In this thresholding task, participants discriminated the direction of the 45°-oriented Gabor patch in the absence of any sounds. Each participant performed 72 trials of the thresholding task and the individual contrast thresholds were used for the main experiment. Prior to performing the thresholding task, participants performed 36 practice trials without a cue.

### EEG Recording and Analysis

Electroencephalogram (EEG) was recorded continuously from 32 Ag/AgCl electrodes mounted in an elastic cap and amplified by an ActiCHamp amplifier (BrainProducts, GmbH). Electrodes were arranged according to the 10-20 system. The horizontal electrooculogram (HEOG) was recorded from two additional electrodes placed on the external ocular canthi which were grounded with an electrode placed on the neck of the participant. The vertical electrooculogram was measured at electrodes FP1 or FP2, located above the left and right eye, respectively. All scalp electrodes were referenced to the right mastoid online and were digitized at 500 Hz.

Data processing was carried out using EEGLAB (Delorme & Makeig, 2004) and ERPLAB (Lopez-Calderon & Luck, 2014) toolboxes and custom-written scripts in MATLAB (The MathWorks, Natick, MA). Continuous EEG data were filtered with a bandpass (butterworth filter) of 0.01-112.5Hz offline. Data were epoched from −1,000 ms to +1,200 ms with respect to the onset of the auditory cue. Artifacts were detected in the time window −800 to 800 ms, and trials contaminated with blinks, eye movements, or muscle movements were removed from the analysis. First, we used automated procedures implemented in ERPLAB (Lopez-Calderon & Luck, 2014; peak-to-peak for blinks, and a step function to detect horizontal eye movements at the HEOG channel). Second, for each participant, each epoch was visually inspected to check the automated procedure and the trials chosen for rejection were updated (cf., Störmer, Alvarez, & Cavanagh, 2014). Artifact-free data was digitally re-referenced to the average of the left and right mastoids. In order to avoid overlap of the target-elicited neural activity with the cue-elicited neural activity, only trials without a target and trials with a 630 ms cue-target SOA were included in the cue-elicited ERP analysis.

ERPs elicited by the left and right noise bursts were averaged separately and were then collapsed across cue position (left, right) and hemisphere of recording (left, right) to obtain waveforms recorded ipsilaterally and contralaterally relative to the sound. The ERPs elicited by the central cues were obtained by averaging across the same lateral electrode positions across both hemispheres (bilateral) that were included in the peripheral cue analysis. ERPs were low-pass filtered (half-amplitude cutoff at 25 Hz; slope of 12dB/octave) to remove high-frequency noise. Mean amplitudes for each participant and condition were measured with respect to a 200 ms prestimulus period (-200 to 0 ms from cue onset), and mean amplitudes were statistically compared using both repeated-measures Analyses of Variance (ANOVAs) and planned follow-up paired t-tests. Our main analysis was focused on the ERP activity during the Auditory-Evoked Contralateral Occipital Positivity (ACOP) time window. The ACOP – usually measured as a contralateral-vs.-ipsilateral ERP component – has been proposed as an index of exogenous attention (McDonald et al., 2013). Thus, based on previous studies on the ACOP (McDonald et al., 2013; Keefe & Störmer, 2020), the ERP amplitude was measured between 260-360 ms at four parietal-occipital electrode sites (PO7/PO8/P7/P8) separately for the hemisphere contralateral to the cued location, ipsilateral to the cued location, and over bilateral electrode sites following the central cue. A separate and more exploratory (though preregistered) analysis focused on frontal activity related to shifting vs. not shifting attention to peripheral and central cues respectively. Based upon prior research demonstrating shift-related activity at frontal sites in endogenous attentional cueing paradigms (Störmer et al., 2009), activity was measured between 300-500 ms at four frontal electrode sites (F3/F4/FC1/FC2).

### Topographical maps

To illustrate the scalp distribution of the different ERP measures, we created topographical maps using spline interpolation of the voltage differences between the cue conditions. To isolate the activity related to the three different cues, we created maps for the contralateral-minus-ipsilateral activity (the ACOP), as well as for the voltage differences between each lateralized activity and the non-lateralized activity elicited by the central cue (i.e., contralateral-minus-central; ipsilateral-minus-central). For contralateral-minus-ipsilateral topographies, values at midline electrode sites (e.g., POz) were set to zero (Störmer et al., 2009), and these difference voltage topographies were projected to the right side of the head. The contralateral/ipsilateral-minus-central topographies were plotted together, with differences in the ipsilateral hemisphere projected to the left side of head and differences in the contralateral hemisphere projected to the right side.

### Statistical Analyses

Behavior was analyzed by comparing accuracy (% correct) in the Gabor discrimination task separately for when a cue was presented at the same location as the visual target (valid trials) vs. at the opposite location (invalid trials) vs. at the center (central trials). Though the behavioral measure of interest was accuracy, we also analyzed reaction time (i.e., RT) in order to rule out any speed-accuracy trade-offs. Behavioral and EEG data were statistically analyzed using repeated-measures ANOVAs and paired t-tests (alpha = 0.05) using MATLAB (The MathWorks, Natick, MA). To compare accuracy and RT in each task following the different cue conditions, we performed 3 × 2 repeated-measures ANOVAs with factors of cue type (valid, invalid, or central) and cue-target SOA (130 ms or 630 ms). Note that the inclusion of the cue-target SOA factor is a departure from our pre-registered analysis but was necessary in order to compare the strength of the alerting response elicited by each cue. To compare ERP activity following each cue, we performed repeated-measures ANOVAs with a factor of hemisphere relative to cue type (contralateral to peripheral cue, ipsilateral to peripheral cue, bilateral for central cue) on the data separately for each of our a priori chosen electrode clusters and time window pairs. Both pre-registered and post-hoc t-tests were performed on the data and are appropriately noted in the Results section. Post-hoc t-tests were corrected for multiple comparisons using a Holm-Bonferroni correction (Holm, 1979) and reported in corrected form.

An additional time-frequency analysis of lateralized and nonlateralized oscillatory activity in the alpha-band (8-13Hz) was pre-registered. These data are not reported here due to the critical a priori ANOVA comparing alpha-frequency activity across the three cue conditions (contralateral vs. ipsilateral vs. central) failing to reach statistical significance. However, the method and results of this analysis are outlined in the Supplementary Alpha Analysis Method and Supplementary Alpha Analysis Results.

## Results

### Behavior

Accuracy following each cue and cue-target SOA in the target discrimination task is plotted in Figure 1B. In order to test for the presence of a behavioral cueing benefit at each SOA, a two-way repeated-measures ANOVA with factors of cue condition (valid, invalid, central) and SOA (short, long) was performed. This analysis revealed a significant main effect of SOA, *F*(1, 18) = 36.00, *p* < .001, η_p_^2^ = 0.59, indicating that overall accuracy was higher in short SOA trials than long SOA trials. Additionally, there was a main effect of cue condition, *F*(2, 36) = 7.79, *p* = 0.002, η_p_^2^ = 0.33, indicating that accuracy varied by cue condition. Finally, there was trend towards a significant interaction between cue condition and SOA, *F*(2, 36) = 3.08, *p* = 0.06, η_p_^2^ = 0.15. Preregistered follow-up t-tests were performed for the short SOA condition, revealing that accuracy was significantly higher following valid cues compared to invalid cues, *t*(18) = 4.39, *p* < 0.001, *d* = 1.01, and central cues, *t*(18) = 3.67, *p* = 0.002, *d* = 0.84. Critically, there was no significant difference between performance following invalid and central cues, *t*(18) = 1.69, *p* = 0.11, *d* = 0.39. This pattern of findings is in line with the predictions of a facilitation-only account of exogenous attention. Post-hoc t-tests were also performed on the long SOA data in order to test for the presence of cueing effects, which were not predicted given that exogenous attention effects are typically largest at short SOAs (Nakayama & Mackeben, 1989). These t-tests, which were corrected for multiple comparisons, demonstrated that accuracy was comparable across all cue conditions. Accuracy was not significantly different following valid and invalid cues, *t*(18) = 1.53, *p* = 0.52, *d* = 0.35, or central cues, *t*(18) = 0.25, *p* = 0.81, *d* = 0.06. Additionally, performance did not significantly differ following central and invalid cues, *t*(18) = 1.20, *p* = 0.50, *d* = 0.28.

**Figure 1.**
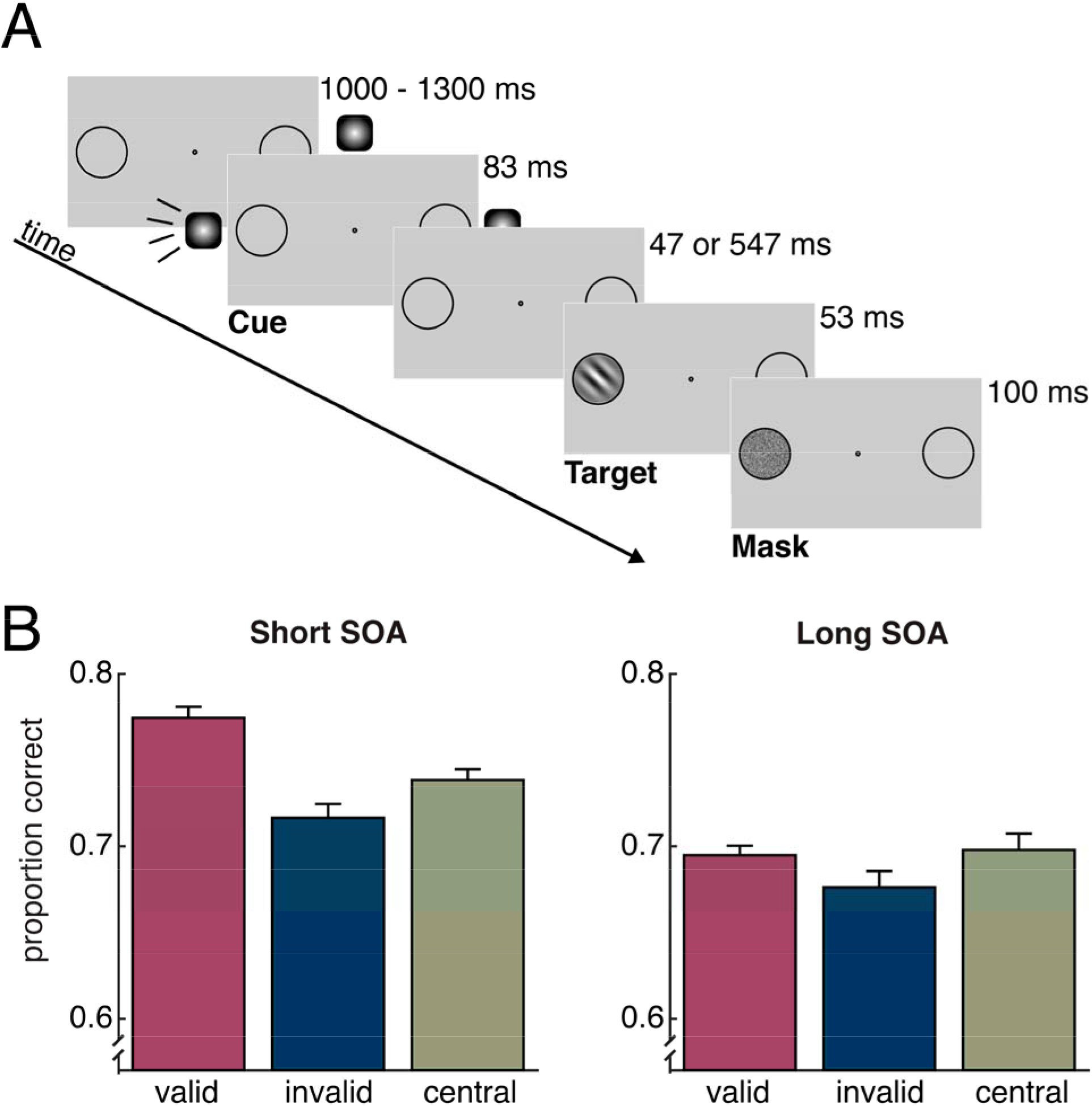
Example trial and performance. (A) Participants discriminated the direction of rotation (clockwise or counterclockwise) of a masked Gabor patch target. Prior to the appearance of the target, participants were presented with an auditory cue that was presented randomly either 130 ms or 630 ms prior to the target. This sound was a pink noise burst played from either the left, right, or center of the screen. On one third of the trials, no visual target was presented. (B) Target discrimination accuracy, plotted as a function of cue condition for each of the cue-target SOAs. Error bars represent ± 1 standard error of the mean.

In order to confirm that any differences in accuracy were not the result of a speed-accuracy trade-off, we analyzed reaction times (i.e. RTs) to the target, plotted in Supplementary Figure 1. We tested whether there were any differences in RT between our conditions of interest by performing a two-way repeated-measures ANOVA with factors of cue validity (valid, invalid, central cue) and SOA (short vs. long) on the RT data. Neither the main effect of SOA, *F*(1, 18) = 3.20, *p* = 0.09, η_p_^2^ = 0.08, nor the main effect of cue condition, *F*(2, 36) = 1.23, *p* = 0.30, η_p_^2^ = 0.08, reached significance. Additionally, there was no significant interaction between cue condition and SOA, *F*(2, 36) = 1.40, *p* = 0.26, η_p_^2^ = 0.07. These findings demonstrate that higher accuracy following the valid vs. invalid and central cues at the short SOA cannot be explained by a trade-off between speed and accuracy.

### Frontal ERPs

Previous research has demonstrated that slow positive deflections in the ERP emerge bilaterally over frontal areas following endogenous symbolic cues that prompt a shift of attention to peripheral locations vs. symbolic cues that do not prompt a shift of attention, termed the Shift-Related Positivity (SRP; Störmer et al., 2009). Based upon this finding, we investigated whether a similar ERP signature associated with the spatial shifting of attention emerged following the random peripheral (shift) vs. central (no-shift) cues over the same time interval in the present study – which would provide support for the idea that our central and peripheral cues oriented attention differently and would also demonstrate the presence of a novel frontal ERP signature of exogenous attentional deployment to uninformative, peripheral auditory cues.

As can be seen in Figure 2, we found a sustained bilateral positivity at frontal sites following the shift vs. no-shift cues. A one-way repeated-measures ANOVA with a factor of cue condition (contralateral to cued location, ipsilateral to cued location, bilateral for central cue) was performed on the ERP waveforms during the predefined SRP time window (300 – 500 ms post-cue). This analysis revealed a main effect of cue condition, *F*(2, 36) = 14.67, *p* < 0.001, η_p_^2^ = 0.45, indicating a significant difference between the amplitude of the waveforms. Planned follow-up t-tests indicated that both the ipsilateral waveform, *t*(18) = 4.03, *p* < .001, *d* = 0.93, and contralateral waveform, *t*(18) = 3.95, *p* < .001, *d* = 0.91, were significantly more positive than the waveform elicited by the central cue. Critically, there was no significant difference between the amplitude of the ipsilateral and contralateral waveforms, *t*(18) = 0.90, *p* = 0.38, *d* = 0.21, indicating that this bilateral frontal component was not sensitive to the specific peripheral location attention was shifted to, but instead indexes control processes related to the spatial orienting response more generally.

**Figure 2.**
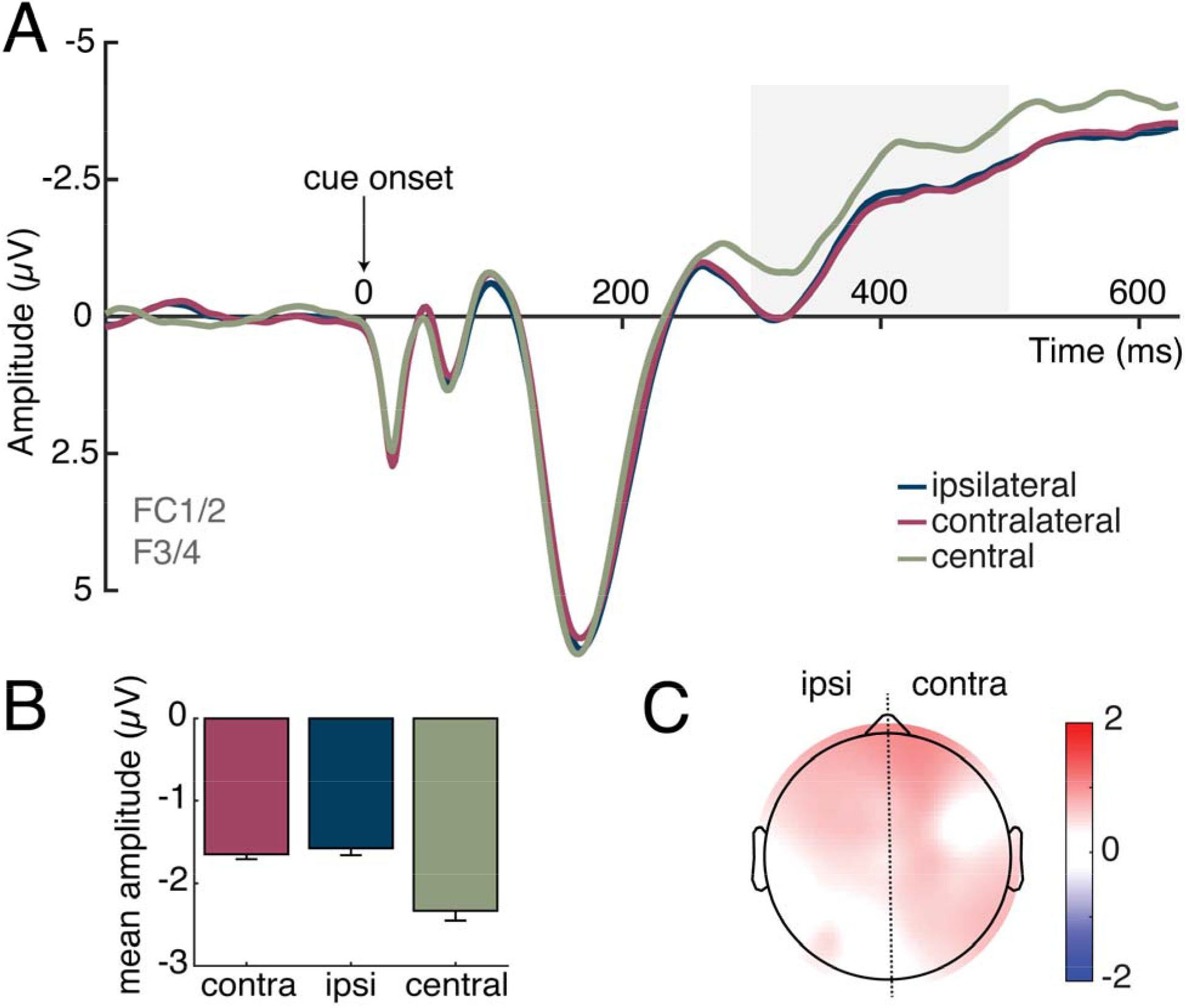
Grand-average ERP waveforms, mean amplitudes, and ERP topography during the shift-related potential (SRP) time window. ERPs at frontal scalp sites (F3/F4/FC1/FC2) were collapsed over left- and right-cue conditions and left and right hemispheres to obtain waveforms recorded ipsilaterally and contralaterally to the cued location. For the central-cue condition, ERPs were computed by averaging bilaterally across the same set of electrodes. (A) Plot of cue-elicited ERP. There was a significant positivity in ipsilateral and contralateral hemispheres in comparison to the no-shift cue (central) during the a priori defined SRP time window (highlighted in gray). (B) Plot of average ERP magnitude during the a priori SRP time window (300-500ms). There was a positivity in ipsilateral (ipsi) and contralateral (contra) cortex relative to the central cue. Error bars represent ± 1 standard error of the mean. (C) Topographical voltage map of the contralateral- and ipsilateral-central ERP difference amplitudes during the SRP time window, with contralateral and ipsilateral differences projected to the right and left sides, respectively. The map demonstrates that the SRP was broadly distributed over bilateral frontal areas, concurrent with the posterior lateralized positivity.

### Occipital ERPs

In order to investigate whether cross-modal exogenous attention improved performance on the visual task by facilitating visual processing at the cued location, suppressing visual processing at the uncued location, or both, we examined the ERPs elicited by the peripheral cues relative to the central cues at parietal-occipital electrode sites. In particular, we focused on the time window of the ACOP – an ERP component that has previously been associated with the exogenous orienting of attention (McDonald et al., 2013; Keefe & Störmer, 2020). As this component is based on a relative difference between neural activity across the hemispheres, it is unclear whether it reflects a contralateral positivity, which would be consistent with visual-cortical enhancement of the attended location; or an ipsilateral negativity, which would be consistent with visual-cortical suppression of the unattended location; or both. If the ACOP reflects facilitation of information at the cued location, then we would expect the contralateral waveform (which reflects activity at the cued location) to be significantly greater in amplitude than the central-cue waveform. In this case, activity ipsilateral to the cue (which reflects activity at the uncued location) should be roughly equivalent to the central-cue baseline. Conversely, if the ACOP reflects suppression of unattended information, then we would expect the ipsilateral waveform to be significantly lower in amplitude than the central-cue waveform. In this case, the contralateral waveform should be comparable to the central-cue baseline. Finally, if the ACOP reflects both facilitation and suppression, then we would expect to observe both a contralateral increase and ipsilateral decrease relative to the central-cue baseline.

As shown in Figure 3, the contralateral ERP waveform was more positive than both the waveform elicited over the ipsilateral hemisphere relative to the peripheral cue as well as the waveform elicited by the central cue over bilateral sites, while there was no difference between ipsilateral and central-cue waveforms. A one-way repeated-measures ANOVA with a factor of cue condition (contralateral to cued location, ipsilateral to cued location, bilateral for central cue) was performed on the ERP waveforms during the ACOP time window (260 – 360 ms post-cue). This analysis revealed a main effect of cue condition, *F*(2, 36) = 11.91, *p* < 0.001, η_p_^2^ = 0.40, indicating a significant difference between the amplitude of the waveforms. Planned follow-up t-tests indicated that the contralateral waveform was significantly more positive than both the ipsilateral, *t*(18) = 4.39, *p* < .001, *d* = 1.01, and central-cue waveform, *t*(18) = 4.57, *p* < .001, *d* = 1.05. There was no significant difference between the amplitude of the ipsilateral and central-cue waveforms, *t*(18) = 0.76, *p* = 0.46, *d* = 0.18.

**Figure 3.**
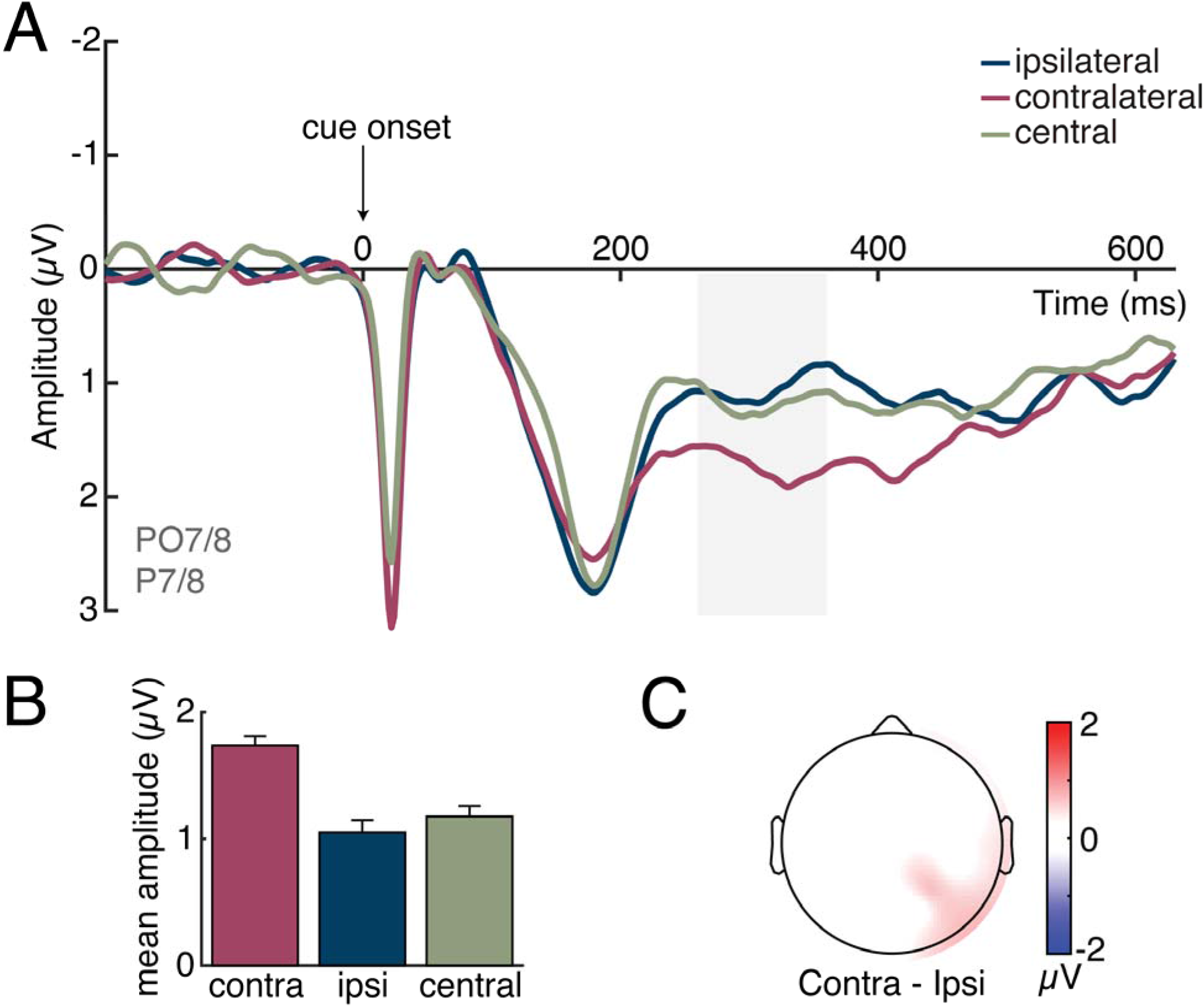
Grand-average ERP waveforms, mean amplitudes, and ERP topography during the ACOP time window. For peripheral cues, ERPs were collapsed over left- and right-cue conditions and left and right hemispheres to obtain waveforms recorded ipsilaterally and contralaterally to the cued location. For the central-cue condition, ERPs were computed by averaging bilaterally across the same set of electrodes. (A) Plot of cue-elicited ERP. There was a significant positivity over the contralateral hemisphere in comparison to the no-shift cue baseline during the a priori defined ACOP time window (highlighted in gray), whereas the ipsilateral waveform was roughly equivalent to the baseline. (B) Plot of average waveform magnitude during the a priori ACOP time window (260-360ms). There was a positivity over contralateral (contra) cortex relative to ipsilateral (ipsi) cortex and relative to the central cue. Error bars represent ± 1 standard error of the mean. (C) Topographical voltage map of the contralateral-minus-ipsilateral ERP difference amplitudes, projected to the right side of the scalp during the ACOP time window. This map demonstrates that the topography of the ACOP was distributed over parietal and occipital areas, with no evidence of lateralized frontal activity.

## Discussion

How does exogenous attention improve visual perception? While several studies have looked at how endogenous attentional orienting affects visual-cortical processing prior to the onset of a target, it is largely unknown how an exogenous attention cue modulates visual-cortical responses. To fill this gap, we recorded EEG activity while subjects performed a cross-modal, exogenous cueing task in which either peripheral left and right sounds or a central, no-shift sound preceded a masked visual target. By including the central cue condition that did not evoke lateral shifts of attention, we were able to examine neural changes triggered by the peripheral cues relative to this baseline condition to differentiate between spatially specific increases and decreases in neural activity and behavioral performance. We found that accuracy was higher following valid vs. no-shift and invalid cues – and that this effect of cue validity was accompanied by an increase in neural activity over parieto-occipital cortex in the hemisphere contralateral to the cue location, relative to the ipsilateral hemisphere as well as the no-shift cue condition. These results indicate that exogenous orienting of spatial attention results in visual-cortical facilitation in the hemisphere contralateral to the attended location, with no signs of suppression in the opposite hemisphere.

Several previous studies have used no-shift cues in behavioral paradigms with the goal of isolating costs and benefits during attentional orienting, with some studies finding costs, others benefits, and some both (e.g., Yeshurun & Carrasco, 1998; Montagna et al., 2009). We believe that the discrepancies between studies are likely due to differences in the exact stimuli and response measures used. For example, other studies have used dependent measures such as RT (Quinlan & Bailey, 1995), gap size threshold in a Landolt square spatial acuity task (Montagna et al., 2009), and d’ (Hermann et al., 2010) or contrast threshold (Pestilli & Carrasco, 2005) in a variable-contrast orientation discrimination task. As behavioral responses are the accumulation of multiple sensory and cognitive processing steps, and different measures may vary in their sensitivity to these processes, it is difficult to pinpoint at what stage costs and benefits arise in these behavioral studies. Using neural measures, as done here, allows for direct assessment of costs and benefits prior to a behavioral response, demonstrating clearly that early visual-cortical processing is enhanced at the attended location. Furthermore, the type of cue used as a neutral baseline condition may also play a role. The central sound used in our study was matched to the peripheral cues with regards to its low-level features (i.e., amplitude, frequency spectrum) and provided the same temporal — but not spatial — information about the visual targets. This means that the central and peripheral cues presumably had equivalent processing demands, served as equivalent alerting signals, and required roughly equivalent encoding times (Jonides & Mack, 1984). This is supported by our behavioral data in which we see a decrement of similar magnitude in each cue condition at the long vs. short SOA, indicating that there are no large differences in the (non-spatial) alerting signal triggered by the central and peripheral cues. Furthermore, the frontal ERP index of attentional shifting showed significantly different neural activity for peripheral relative to central cues. Thus, we think the central cue in our study acts as a valid baseline relative to the peripheral cues, as it differs only in the main dimension of interest: shifting attention to a new location vs. not.

In the present study we used auditory cues to orient spatial attention, which allowed us to examine cue-related activity over occipital cortex without the contamination of visual-evoked responses that are necessarily triggered when using visual peripheral cues. Thus, the present paradigm extends previous work on visual attentional cueing, demonstrating that exogenous attention effectively operates across modalities, in support of a supramodal account of attention (Farah et al., 1989). It seems particularly important to understand how spatial attention operates cross-modally, as in the real world our attention is often captured by auditory and not only visual stimuli (Störmer, 2019). However, one question that may arise is whether our cross-modal findings can be generalized to intra-modal studies of attention. Critically, several cross-modal attention paradigms have found effects similar to visual-only attention paradigms (for a review, see Hillyard et al., 2016), and a recent experiment directly comparing ERP activity triggered by visual and auditory cues found similar lateralized biasing signals over parietal-occipital cortex as here (Störmer et al, 2019), suggesting that the present results would hold for visual cues as well.

The present findings implicate a difference in the neural mechanisms engaged by exogenous and endogenous attention, as prior research has shown that endogenous attention results in both costs and benefits in behavior (Hawkins et al., 1990; Golob, Pratt, & Starr, 2002) and facilitation and suppression of neural activity (Luck et al., 1994; Mangun & Buck, 1996; Schröger & Eimer, 1997). That is, the present results indicate that exogenous attention only facilitates processing of information at the location of a salient cue, whereas endogenous attention seems to additionally involve suppressive attentional mechanisms that possibly emerge later in time. This facilitation-only account of exogenous attention comports well with other recent evidence demonstrating that lateralized occipital activity predicts behavioral performance only on validly but not invalidly cued trials (Feng et al., 2014; Feng et al., 2017), suggesting that only neural activity at the location of a cue (presumably reflecting the facilitative effects of attention) is related to behavioral performance exogenous cueing tasks. However, it is important to note that the present study only investigated the first ~600 ms of cue-elicited neural activity, in line with the rapid time course of exogenous attention. Accordingly, it is possible that there is a later, suppressive component of attention that only emerges at longer timescales – and that thus would likely only be engaged during endogenous attention due to its relatively sluggish response. Indeed, prior research measuring alpha frequency oscillatory activity following informative central cues suggests that suppression may emerge on a later timescale than facilitation (Green & McDonald, 2010). It is worth noting that we did not observe significant lateralized alpha activity in the present study, however, possibly suggesting that separate neural measures (e.g. ERPs and oscillatory activity) may reveal different patterns of hemisphere-specific neural processing in response to peripheral cues. Regardless, the present data also demonstrate a novel similarity between the exogenous and endogenous orienting of attention, as we find that exogenous cues elicit a Shift-Related Positivity over frontal areas that has previously only been demonstrated in response to endogenous cues (Störmer et al., 2009). Taken together, these findings suggest that the shifting of exogenous and endogenous attention may be mediated by similar control processes in frontal areas, but that these shifts result in different effects in posterior cortex.

Overall, our data demonstrate that the exogenous orienting of spatial attention results in visual-cortical enhancement at the location of a salient cue but does not result in spatially specific suppression of visual processing at uncued locations (i.e., opposite hemisphere). Broadly, these findings suggest that exogenous and endogenous spatial attention differ in how they bias visual processing in anticipation of a target to support effective stimulus selection.

## Acknowledgements

We thank Raymond Hallmen III for help with data collection.

## References

Brainard, D. H. (1997). The psychophysics toolbox. Spatial Vision, 10(4), 433–436.

Carrasco, M. (2011). Visual attention: The past 25 years. Vision Research, 51(13), 1484–1525. https://doi.org/10.1016/j.visres.2011.04.012

Carrasco, M., Penpeci-Talgar, C., & Eckstein, M. (2000). Spatial covert attention increases contrast sensitivity across the CSF: Support for signal enhancement. Vision Research, 40(10–12), 1203–1215. https://doi.org/10.1016/S0042-6989(00)00024-9

Couperus, J. W., & Mangun, G. R. (2010). Signal enhancement and suppression during visual–spatial selective attention. Brain Research, 1359, 155–177. https://doi.org/10.1016/j.brainres.2010.08.076

Cutrone, E. K., Heeger, D. J., & Carrasco, M. (2014). Attention enhances contrast appearance via increased input baseline of neural responses. Journal of Vision, 14(14), 16–16. https://doi.org/10.1167/14.14.16

Delorme, A., & Makeig, S. (2004). EEGLAB: An open source toolbox for analysis of single-trial EEG dynamics including independent component analysis. Journal of Neuroscience Methods, 134(1), 9–21.

Desimone, R., & Duncan, J. (1995). Neural mechanisms of selective visual attention. Annual Review of Neuroscience, 18(1), 193–222.

Farah MJ, Wong AB, Monheit MA, Morrow LA (1989) Parietal lobe mechanisms of spatial attention - modality-specific or supramodal. Neuropsychologia 27:461–470.

Feng, W., Stormer, V. S., Martinez, A., McDonald, J. J., & Hillyard, S. A. (2014). Sounds Activate Visual Cortex and Improve Visual Discrimination. Journal of Neuroscience, 34(29), 9817–9824. https://doi.org/10.1523/JNEUROSCI.4869-13.2014

Feng, W., Störmer, V. S., Martinez, A., McDonald, J. J., & Hillyard, S. A. (2017). Involuntary orienting of attention to a sound desynchronizes the occipital alpha rhythm and improves visual perception. Neuroimage, 150, 318–328.

Golob, E. (2002). Preparatory slow potentials and event-related potentials in an auditory cued attention task. Clinical Neurophysiology, 113(10), 1544–1557. https://doi.org/10.1016/S1388-2457(02)00220-1

Green, J. J., & McDonald, J. J. (2010). The role of temporal predictability in the anticipatory biasing of sensory cortex during visuospatial shifts of attention. Psychophysiology, 47(6), 1057–1065.

Grent-‘t-Jong, T., & Woldorff, M. G. (2007). Timing and Sequence of Brain Activity in Top-Down Control of Visual-Spatial Attention. PLOS Biology, 5(1), e12. https://doi.org/10.1371/journal.pbio.0050012

Hawkins, H. L., Hillyard, S. A., Luck, S. J., Mouloua, M., Downing, C. J., & Woodward, D. P. (1990). Visual attention modulates signal detectability. Journal of Experimental Psychology: Human Perception and Performance, 16(4), 802–811. https://doi.org/10.1037/0096-1523.16.4.802

Herrmann, K., Montaser-Kouhsari, L., Carrasco, M., & Heeger, D. J. (2010). When size matters: Attention affects performance by contrast or response gain. Nature Neuroscience, 13(12), 1554–1559. https://doi.org/10.1038/nn.2669

Holm, S. (1979). A Simple Sequentially Rejective Multiple Test Procedure. Scandinavian Journal of Statistics, 6(2), 65–70. JSTOR.

Jonides, J., & Mack, R. (n.d.). On the Cost and Benefit of Cost and Benefit. 16.

Keefe, J. M., & Störmer, V. S. (2020). Alpha-band oscillations and slow potentials shifts over visual cortex track the time course of both endogenous and exogenous orienting of attention. BioRxiv, 2019.12.12.874818. https://doi.org/10.1101/2019.12.12.874818

Kelly, S. P., Lalor, E. C., Reilly, R. B., & Foxe, J. J. (2006). Increases in alpha oscillatory power reflect an active retinotopic mechanism for distracter suppression during sustained visuospatial attention. Journal of neurophysiology, 95(6), 3844–3851.

Lopez-Calderon, J., & Luck, S. J. (2014). ERPLAB: An open-source toolbox for the analysis of event-related potentials. Frontiers in Human Neuroscience, 8, 213.

Luck, S. J., Hillyard, S. A., Mouloua, M., Woldorff, M. G., Clark, V. P., & Hawkins, H. L. (n.d.). Effects of Spatial Cuing on Luminance Detectability: Psychophysical and Electrophysiological Evidence for Early Selection. 18.

Mangun, G. R., & Buck, L. A. (1998). Sustained visual-spatial attention produces costs and benefits in response time and evoked neural activity. Neuropsychologia, 36(3), 189–200. https://doi.org/10.1016/S0028-3932(97)00123-1

McDonald, J. J., Stormer, V. S., Martinez, A., Feng, W., & Hillyard, S. A. (2013). Salient Sounds Activate Human Visual Cortex Automatically. Journal of Neuroscience, 33(21), 9194–9201. https://doi.org/10.1523/JNEUROSCI.5902-12.2013

McDonald, John J., & Green, J. J. (2008). Isolating event-related potential components associated with voluntary control of visuo-spatial attention. Brain Research, 1227, 96–109. https://doi.org/10.1016/j.brainres.2008.06.034

McDonald, John J., Teder-Sälejärvi, W. A., & Hillyard, S. A. (2000). Involuntary orienting to sound improves visual perception. Nature, 407(6806), 906–908. https://doi.org/10.1038/35038085

Montagna, B., Pestilli, F., & Carrasco, M. (2009). Attention trades off spatial acuity. Vision Research, 49(7), 735–745. https://doi.org/10.1016/j.visres.2009.02.001

Nakayama, K., & Mackeben, M. (1989). Sustained and transient components of focal visual attention. Vision Research, 29(11), 1631–1647. https://doi.org/10.1016/0042-6989(89)90144-2

Nobre, A. C., Sebestyen, G. N., & Miniussi, C. (2000). The dynamics of shifting visuospatial attention revealed by event-related potentials. Neuropsychologia, 38(7), 964–974. https://doi.org/10.1016/S0028-3932(00)00015-4

Pelli, D. G. (1997). The VideoToolbox software for visual psychophysics: Transforming numbers into movies. Spatial Vision, 10(4), 437–442.

Pestilli, F., & Carrasco, M. (2005). Attention enhances contrast sensitivity at cued and impairs it at uncued locations. Vision Research, 45(14), 1867–1875. https://doi.org/10.1016/j.visres.2005.01.019

Pinsk, M. A., Doniger, G. M., & Kastner, S. (2004). Push-Pull Mechanism of Selective Attention in Human Extrastriate Cortex. Journal of Neurophysiology, 92(1), 622–629. https://doi.org/10.1152/jn.00974.2003

Quinlan, P. T., & Bailey, P. J. (1995). An examination of attentional control in the auditory modality: Further evidence for auditory orienting. Perception & Psychophysics, 57(5), 614–628. https://doi.org/10.3758/BF03213267

Schröger, E., & Eimer, M. (1997). Endogenous Covert Spatial Orienting in Audition Cost-Benefit Analyses of Reaction Times and Event related Potentials. The Quarterly Journal of Experimental Psychology Section A, 50(2), 457–474. https://doi.org/10.1080/713755706

Serences, J. T., Yantis, S., Culberson, A., & Awh, E. (2004). Preparatory Activity in Visual Cortex Indexes Distractor Suppression During Covert Spatial Orienting. Journal of Neurophysiology, 92(6), 3538–3545. https://doi.org/10.1152/jn.00435.2004

Störmer, V. S. (2019). Orienting spatial attention to sounds enhances visual processing. Current Opinion in Psychology, 29, 193–198.

Störmer, V. S., Alvarez, G. A., & Cavanagh, P. (2014). Within-hemifield competition in early visual areas limits the ability to track multiple objects with attention. Journal of Neuroscience, 34(35), 11526–11533.

Störmer, V. S., Green, J. J., & McDonald, J. J. (2009). Tracking the voluntary control of auditory spatial attention with event-related brain potentials. Psychophysiology, 46(2), 357–366.

Störmer, V. S., McDonald, J. J., & Hillyard, S. A. (2009). Cross-modal cueing of attention alters appearance and early cortical processing of visual stimuli. Proceedings of the National Academy of Sciences, 106(52), 22456–22461. https://doi.org/10.1073/pnas.0907573106

Watson, A. B., & Pelli, D. G. (1983). QUEST: A Bayesian adaptive psychometric method. Perception & Psychophysics, 33(2), 113–120.

Yeshurun, Y., & Carrasco, M. (1999). Spatial attention improves performance in spatial resolution tasks. Vision Research, 14.

