## Supplement for "Cross-modal orienting of exogenous attention results in visual-cortical facilitation, not suppression"

Dartmouth College

**Supplementary Alpha Analysis Method**

For the time frequency analysis, single-trial EEG data was analyzed via complex Morlet wavelets before averaging, following the methods of Lakatos et al. (2004) and Torrence and Compo (1998). Spectral amplitudes were calculated via four-cycle wavelets at 60 different frequencies increasing linearly from 2 to 40 Hz separately for each electrode, time point (every 2 ms), attention condition (left, right), and participant. Spectral amplitudes were then averaged across trials separately for each condition and participant, and a mean baseline of −350 to –150 ms from cue onset was subtracted from each time point for each frequency separately (Keefe & Störmer, 2020). Mean spectral amplitudes elicited by the left and right noise bursts were then collapsed across cued location (left, right) and lateral position of the electrode (left, right) to reveal attention-induced modulations ipsilateral and contralateral to the cued location. Oscillatory activity elicited by the central noise bursts was analyzed by averaging across the same lateral electrode positions (left, right) that were included in the peripheral cue analysis. The statistical analysis was focused on alpha-band amplitude modulations over the range of 8 – 13 Hz at parietal-occipital electrode sites (PO7/PO8/P7/P8) and during the same time intervals as the ACOP (260 – 360 ms).

**Supplementary Alpha Analysis Results**

In addition to the ERP analysis, we examined the alpha frequency activity elicited by the peripheral cues relative to the central cues at occipital electrode sites in order to investigate whether cross-modal exogenous attention improved performance on the visual task by facilitating processing at the cued location, suppressing processing at the uncued location, or both. For this analysis, we focused on the same time window used in the ACOP analysis – from 260 to 360 ms post-cue. A one-way repeated-measures ANOVA with a factor of cue condition (contralateral, ipsilateral, central) was performed on the alpha power during the ACOP time window. This analysis indicated that there was not a main effect of cue condition, *F*(2, 36) = 2.30, *p* = 0.12, η_p_^2^ = 0.11, indicating a that there was not a significant difference in the amplitudes of alpha changes. Therefore, the alpha analysis was terminated at this point.

**Supplementary Figure 1**

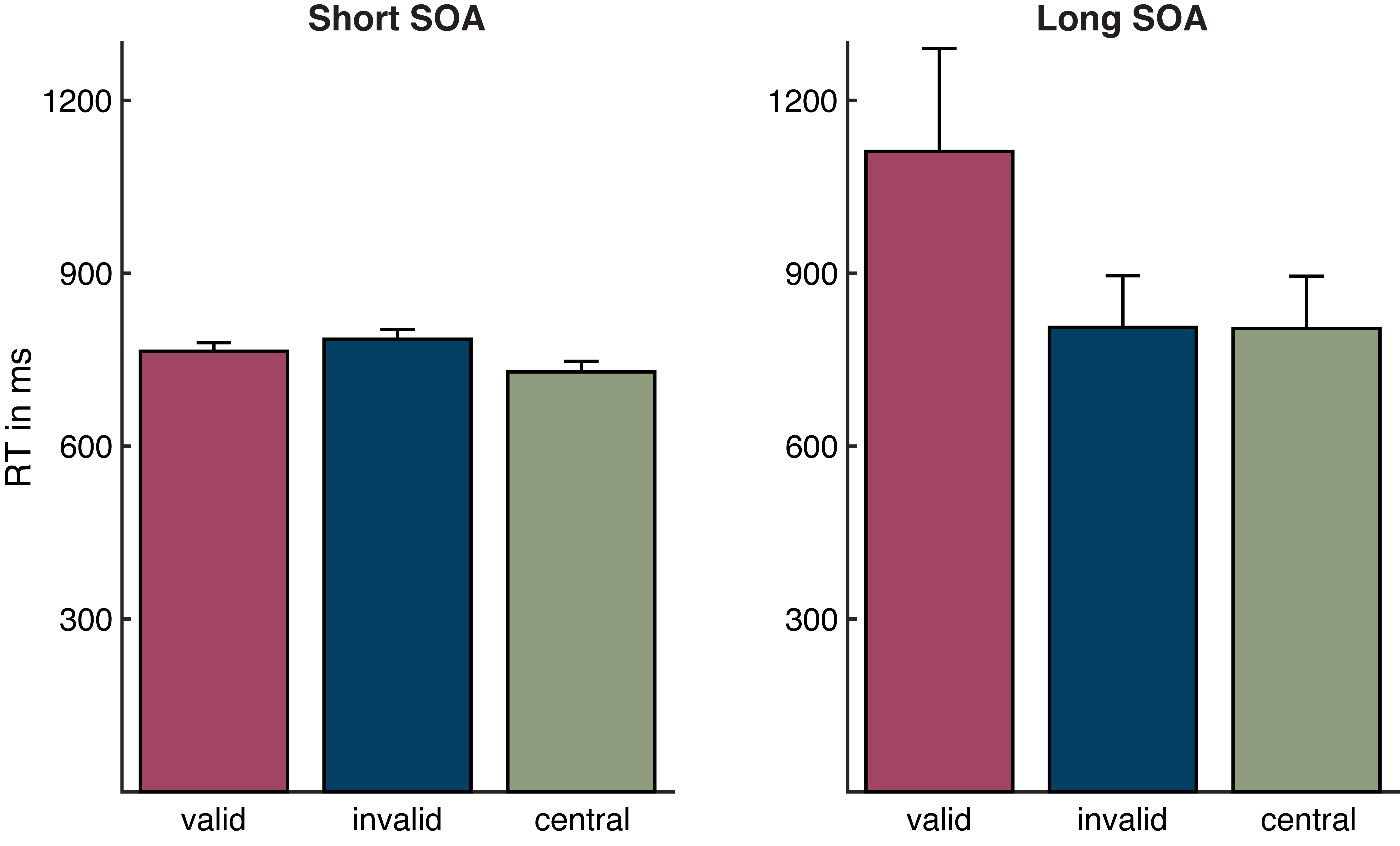

Figure S1. Response time (RT) to the target in each cue condition and stimulus onset asynchrony (SOA) between cue and target. There were no significant differences in RT based upon cue condition nor SOA.
